# The cytolethal distending toxin modulates cell differentiation and elicits epithelial to mesenchymal transition

**DOI:** 10.1101/2022.04.06.487255

**Authors:** Lamia Azzi-Martin, Valentin Touffait-Calvez, Maude Everaert, Ruxue Jia, Elodie Sifré, Lornella Seeneevassen, Christine Varon, Pierre Dubus, Armelle Ménard

## Abstract

We are frequently exposed to bacterial genotoxins, such as cytolethal distending toxin (CDT), a prevalent heterotrimeric toxin whose active moiety is its CdtB subunit. CdtB triggers potent DNA damage, predisposing factors in the development of cancers, in host cells. CDT from *Helicobacter hepaticus*, a mouse pathogen, was shown to be directly involved in the development of murine hepatocarcinoma. Preliminary studies have shown that CDT induces certain phenotypes reminiscent of epithelial to mesenchymal transition (EMT), a process by which cells lose their epithelial characteristics in favor of mesenchymal ones, conducive to cell motility. In the present study, we investigated the different steps of EMT using liver tissues of mice infected with *H. hepaticus*, as well as human epithelial cell lines and xenograft mouse models following *H. hepaticus* CdtB expression. Most of the different steps of the EMT process were reproduced throughout the studied models. Indeed, microarray data showed a CdtB-dependent regulation of EMT-related transcripts. The key transcriptional regulators of EMT (SNAIL1 and ZEB1) and EMT markers (Vimentin, Fibronectin and α5β1 integrin) were upregulated both at RNA and protein levels in response to CdtB. It also induced cell-cell junctions’ disassembly, causing individualization of cells and acquisition of a spindle-like morphology. CdtB activated the expression and activity of matrix metalloproteases and increased cell motility. This study demonstrated that CDT/CdtB elicits EMT process activation, supporting the idea that infection with genotoxin-producing bacteria can promote malignant transformation.

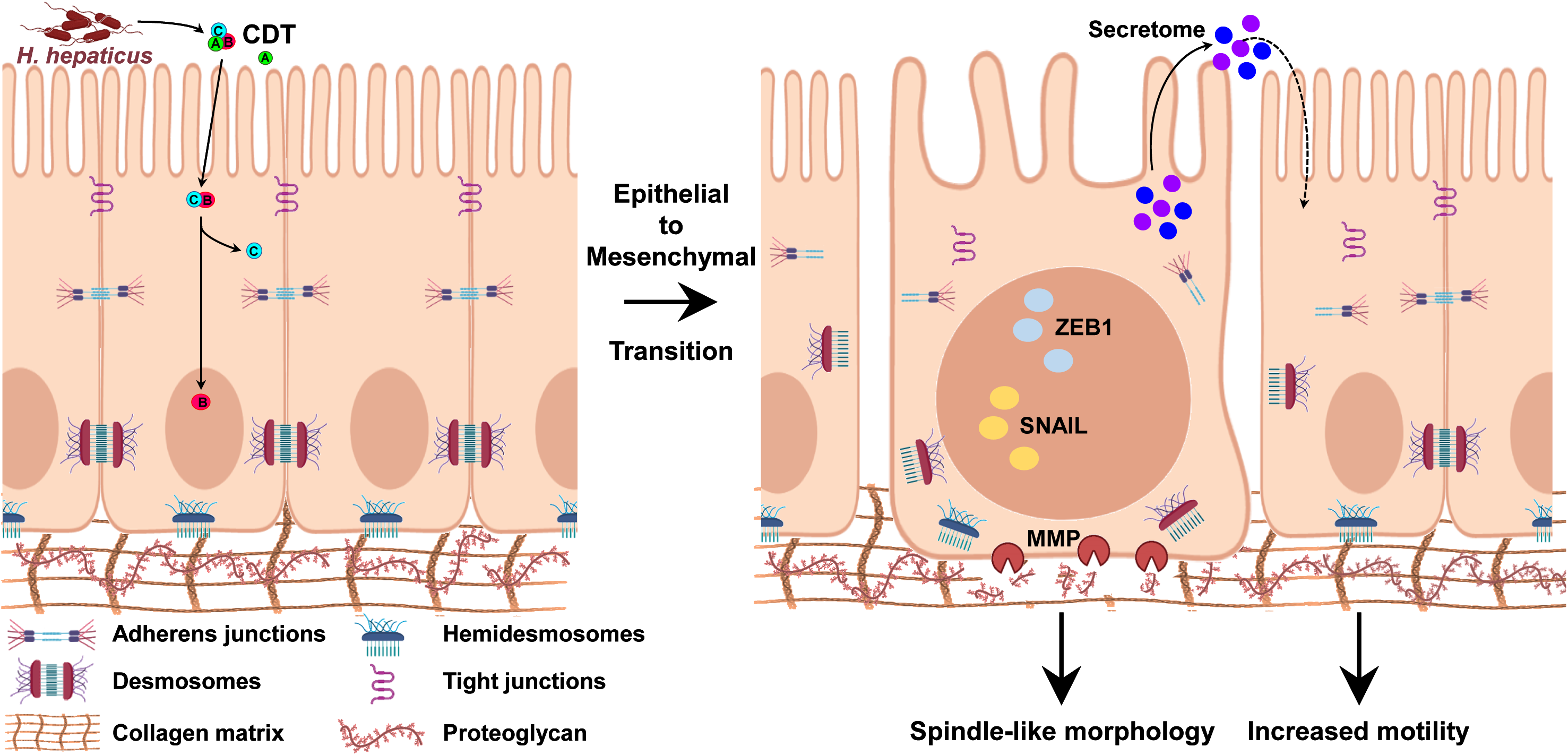

**Author Summary:** We are frequently exposed to infection with genotoxin--producing bacterial from the gut microbiota, such as cytolethal distending toxin (CDT). CDT, *via* its active CdtB subunit, causes severe DNA damage in host cells, well-known risk factor of cancer development and progression. Chronic infection with CDT-producing bacteria is thus involved in cancer development. CDT is widespread among many bacteria and its impact in human cancer seems likely underestimated. Despite its major significance, CDT remains little studied. Here, we showed that cells exposed to CdtB are no longer cohesive, become individualized and acquire a spindle-shaped morphology known to be conducive to migration. These cells also express increased level of mesenchymal markers, as well as increased level of SNAIL1 and ZEB1, two key transcription factors orchestrating a crucial mechanism for cancer initiation and progression: epithelial to mesenchymal transition. These effects induced by CdtB were associated with increased matrix metalloproteinases degrading activity and emergence of cellular motility. Collectively, these data showed that CdB activates epithelial to mesenchymal transition, supporting the role of CDT in tumorigenesis.

## Introduction

Our microbiota composition influences health and disease throughout our life. In this context, bacteria are known to impact cancer initiation, progression and even modulate anticancer drug efficacy. The involvement of chronic bacterial infections in carcinogenesis has been first proven in 1994 following evidence of the role of *Helicobacter pylori* infection in the development of gastric cancers in human [1]. In 1994, a novel *Helicobacter* species, *Helicobacter hepaticus*, was isolated from the liver and colon of mice with active chronic hepatitis [2]. *H. hepaticus* is usually present in mouse intestine. It colonizes the biliary tract and the liver and induces chronic hepatitis in mice, which can lead to hepatocarcinoma in older animals [2]. In these mice infected by *H. hepaticus*, CDT-induced hepatocarcinoma occurs through the Cytolethal Distending Toxin (CDT) [3]. More recently, it has been shown that CDT from *H. hepaticus* also induces colitis and intestinal carcinogenesis in susceptible mice [4]. Several CDT-secreting enterohepatic *Helicobacter* have been found in the bile ducts of patients with cholangiocarcinoma or in the liver of patients with hepatocellular carcinoma, suggesting a role for these bacteria and their toxin in the hepatobiliary carcinogenic process(es) [5, 6].

CDT is widespread among many pathogenic Gram-negative bacteria with an impact on human disease [7]. The involvement of CDT in malignant transformation was demonstrated in several animal models [4, 8]. CDT is a subclass of the AB toxin superfamily and is active as a heterotrimeric complex, composed of three distinct subunits among which CdtB is the active one. CdtB displays dual DNase and phosphatase activities. CdtB triggers DNA damage in host cells [9] and impairs DNA-damage response, leading to genomic instability and accumulation of mutations transmitted to daughter cells [10, 11]. Additionally, CDT induces inflammatory signatures [12, 13], leads to anchorage-independent growth [10] and targets pathways involved in cell transformation, as mitogen-activated protein kinase [10], phosphoinositide 3-kinase- AKT [14], nuclear factor-κB [15], autophagy [16] amongst others [17]. Thus, various CDT- induced effects are common hallmarks of many cancers and indicative of the toxin’s malignant- transformation potential.

Among the several processes involved in cancer initiation and/or progression, Epithelial to Mesenchymal Transition (EMT) is a crucial one. EMT is a reversible cellular reprogramming mechanism by which cells lose their epithelial characteristics to acquire mesenchymal properties conducive to cell motility and invasion. It was shown that radiotherapies and chemotherapies (cisplatin and 5-fluorouracil for example) aiming at damaging DNA can induce EMT [18, 19]. CDT could thus elicit EMT *via* its DNase activity. Indeed, some phenotypes induced by CDT, such as the profound remodeling of actin cytoskeleton, the accumulation of stress fibers and the formation of lamellipodia [20, 21], are reminiscent of EMT-induced phenotypes, suggesting that CDT could stimulate EMT.

EMT is orchestrated by transcription factors (TFs) which repress epithelial genes, activate mesenchymal ones and control EMT-related signaling pathways [22]. The SNAIL and ZEB family TFs are found among the key regulators of EMT. EMT is also characterized by the loss of epithelial cell-cell junction and polarity. Tight junctions are disrupted by dissolution or relocation of junctional proteins. Indeed, decreased expression of claudins and occludins while relocalisation of zonula occludens 1 (ZO-1) is described during EMT [23]. Tight junctions’ dissociation is accompanied by E-cadherin loss which promotes β-catenin release, resulting in the disassembly of Adherens junctions. Desmosomes and gap junctions are also disrupted during EMT. In addition, actin cytoskeleton remodeling and reorganization of stress fibers also occur [24]; cells acquire a spindle-shaped morphology and develop cellular protrusions (lamellipodia, filopodia and invadopodia) accompanied by increased migratory capacity as well as extracellular matrix (ECM) degrading properties, the latter driven by the expression of matrix metalloproteinases (MMPs) [24].

The present study aimed to evaluate the role of CDT/CdtB in modulating cell differentiation and EMT promotion. Accordingly, microarray-based identification of differentially expressed genes was performed to identify EMT-associated genes whose expression is regulated by CdtB of *H. hepaticus*. Then, the effects of CdtB on EMT-TFs were evaluated. Mesenchymal markers expression were also evaluated, as well as that of proteins implicated in cell-cell junctions, cytoskeleton remodeling, ECM degradation and cellular motility. These investigations were performed on human epithelial cell lines upon ectopic expression of *H. hepaticus* CdtB and its corresponding mutated CdtB (H_265_L) lacking catalytic activity, to attribute the observed effects specifically to the CdtB. Some results were also confirmed using liver tissues of mice infected with *H. hepaticus* [25] and xenograft mouse models [26], as these models have already been validated for CDT/CdtB study.

## Results

It has been reported that *Helicobacter hepaticus* colonizes primarily the intestines and liver, and this infection exacerbates the development of cancer at both hepatic, intestinal, and extraintestinal sites [27]. The effects of *H. hepaticus* infection in triggering EMT were first assessed on infected mice. Then, the effects of CDT/CdtB were assessed on human intestinal and hepatic epithelial cell lines following the cellular ectopic expression of *H. hepaticus* CdtB and its corresponding mutated CdtB (H_265_L) lacking catalytic activity [28] to examine the effects specifically related to the CdtB subunit.

### *Helicobacter hepaticus* infection regulates the expression and localization of EMT markers

Immunohistochemical analyses of the liver of mice infected with *H. hepaticus* for 14 monthspreviously showed that *H. hepaticus* infection triggers liver lesions [25], as well as the nuclear remodeling of hepatocytes [29]. Analysis of E-cadherin staining, an epithelial marker, revealed a 3.5-fold reduced stained area following *H. hepaticus* infection, suggesting the loss of Adherens junctions (Fig. 1A,D). The analysis of ZEB1, ZEB2 and SNAIL EMT TFs in the liver of these mice was not conclusive. However, TWIST analysis revealed a 3.4-fold increase of this EMT TF in the big nuclei of hepatocytes of *H. hepaticus* infected mice, compared to non-infected mice (Fig. 1B,D). Analysis of the mesenchymal marker Vimentin, a type III intermediate filament protein, revealed a 5.2-fold increased stained area following *H. hepaticus* infection. Moreover, the increase of Vimentin staining was observed both for hepatocytes and sinusoidal endothelial cells in the liver of infected mice (Fig. 1C,D) along with an obvious increase in intensity. Together these data showed that *H. hepaticus* regulates the expression and localization of EMT markers in the liver of infected mice, suggesting that *H. hepaticus* infection elicits EMT process. Moreover, obvious TWIST accumulation in the bigger nuclei indicates that CDT of *H. hepaticus* may play a role in the onset of EMT.

**Fig. 1.**
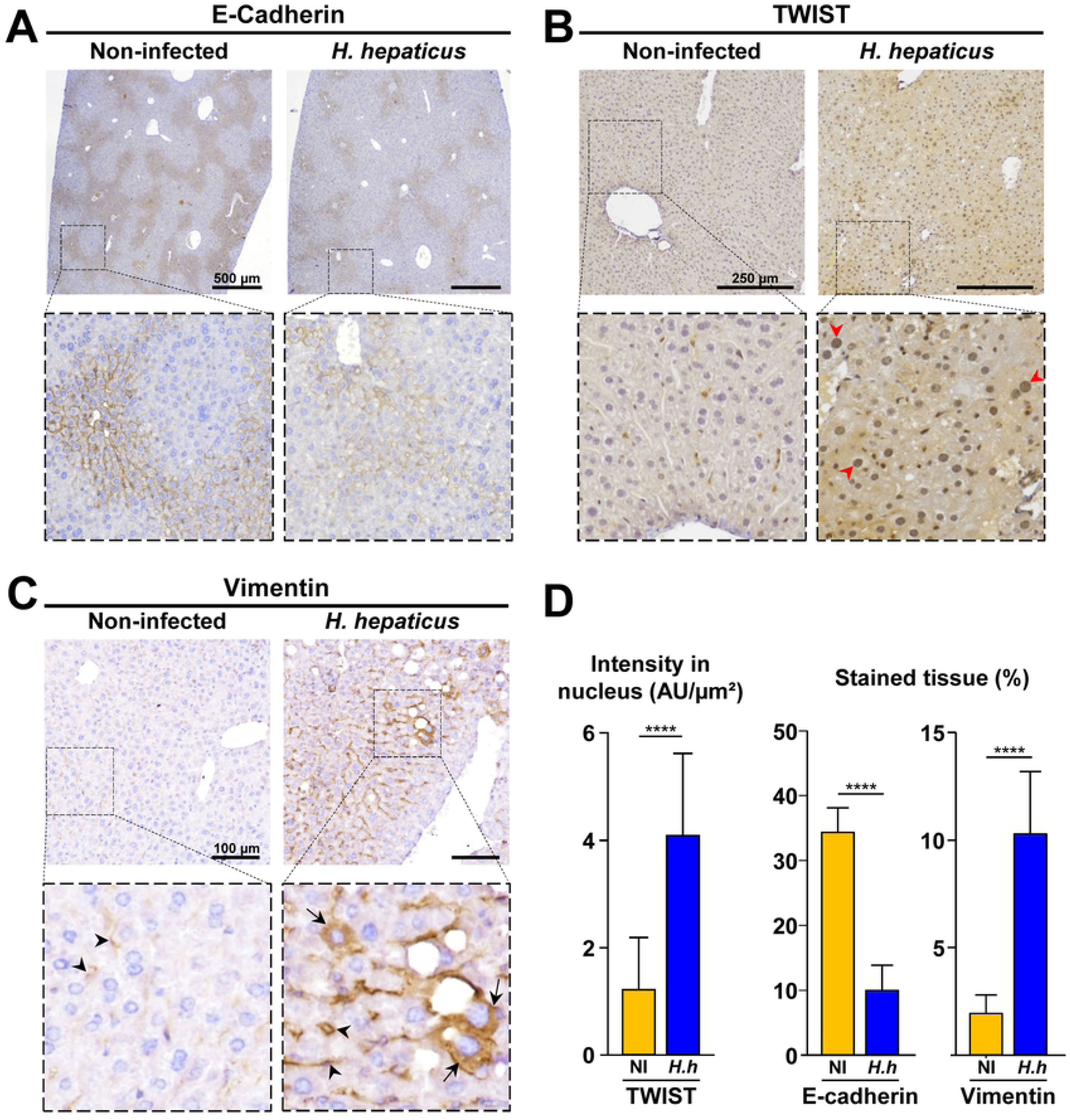
*Helicobacter hepaticus* infection regulates the expression and subcellular localization of EMT protein markers in the liver of infected mice. Non-transgenic mice were infected or not with *H. hepaticus* wild type strain 3B1 for 14 months [25]. Then three μm- tissue sections of liver specimens were immunostained for E-Cadherin **(A)**, TWIST **(B)** and Vimentin **(C)** and counterstained with standard hematoxylin staining. **(A)** to **(C)** Images of mouse livers following a 14-months infection with *H. hepaticus*. Magnifications of selected areas are shown in boxes. Red arrowheads indicate nucleus concentrating TWIST. Black arrows and arrowheads indicate accumulation of vimentin in the cytoplasm of enlarged hepatocytes and liver sinusoidal endothelial cells, respectively. **(D)** Quantification of EMT protein markers. Nuclear TWIST intensity and the percentage of E- cadherin and vimentin-stained tissues. Results are presented as the mean ± SD. ****p<0.0001 *versus* non-infected mice. AU, arbitrary units; *H.h, Helicobacter hepaticus;* NI, non-infected.

### The CdtB of *Helicobacter hepaticus* regulates EMT-associated genes

Whole human genome microarray was previously performed and validated upon CdtB ectopic expression in epithelial intestinal HT-29 cells [13]. In this model, the expected well-known cellular phenotypes induced by CDT/CdtB were observed, i.e. actin cytoskeleton remodeling, cellular and nuclear distension, and cell cycle arrest [13, 21]. At the mRNA level, EMT is characterized by the downregulation of epithelial cytokeratins and upregulation of mesenchymal proteins. In the present study (Fig. 2), microarrays data regarding the expression of epithelial markers were not conclusive as most of epithelial transcripts were not significantly regulated in response to CdtB (S1 Table) and four epithelial genes were found upregulated in response to CdtB: *CDH1* (E-cadherin), *KRT7* and *KRT14* (keratin 7 and 14), *NECTIN4*, and *OCLN* (Occludin). On the other hand, some mesenchymal markers mRNAs were upregulated following CdtB expression, such as *VIM* and *SPARC* encoding Vimentin and Osteonectin. The expression of *LAMA3*, *LAMB3* and *LAMC2* genes, which encode the subunits of Laminin-5 heterotrimer, an EMT activator, is also activated upon CdtB expression. Other mesenchymal markers genes were found activated by CdtB, as well as *ITGB1* encoding Integrin subunit beta 1. The well-known core EMT-TFs [23], *SNAI2*/*SLUG*, *ZEB2* and *TWIST* were not detected at the basal level (S1 Table), while an increase in *SNAI1*/*SNAIL* and ZEB1 transcripts was observed in response to CdtB. Other EMT-associated mRNA were also upregulated, including *CTNNB1*, *CCN1*, *STAT3*, and *TCF3*, encoding β-catenin, β-catenin, Cellular Communication Network Factor 1, Signal Transducer and Activator of Transcription 3 and Transcription Factor 3. Additionally, CdtB triggered the upregulation of mRNAs of EMT signaling molecules known to be activated (*B2M*, *BMP1*, *CALD1*, *EZR*, GJB4, *JAG1, SRC*) encoding β2- Microglobulin, Bone Morphogenetic Protein 1, Caldesmon 1, Ezrin, Gap Junction Protein Beta 4, Jagged-1 (the Notch ligand) and SRC Proto-Oncogene. Taken together, the upregulation of mesenchymal markers transcripts, as well as that of EMT-TFs, major EMT players and EMT signaling molecules suggests the activation of EMT signaling in response to CdtB intoxication.

**Fig. 2.**
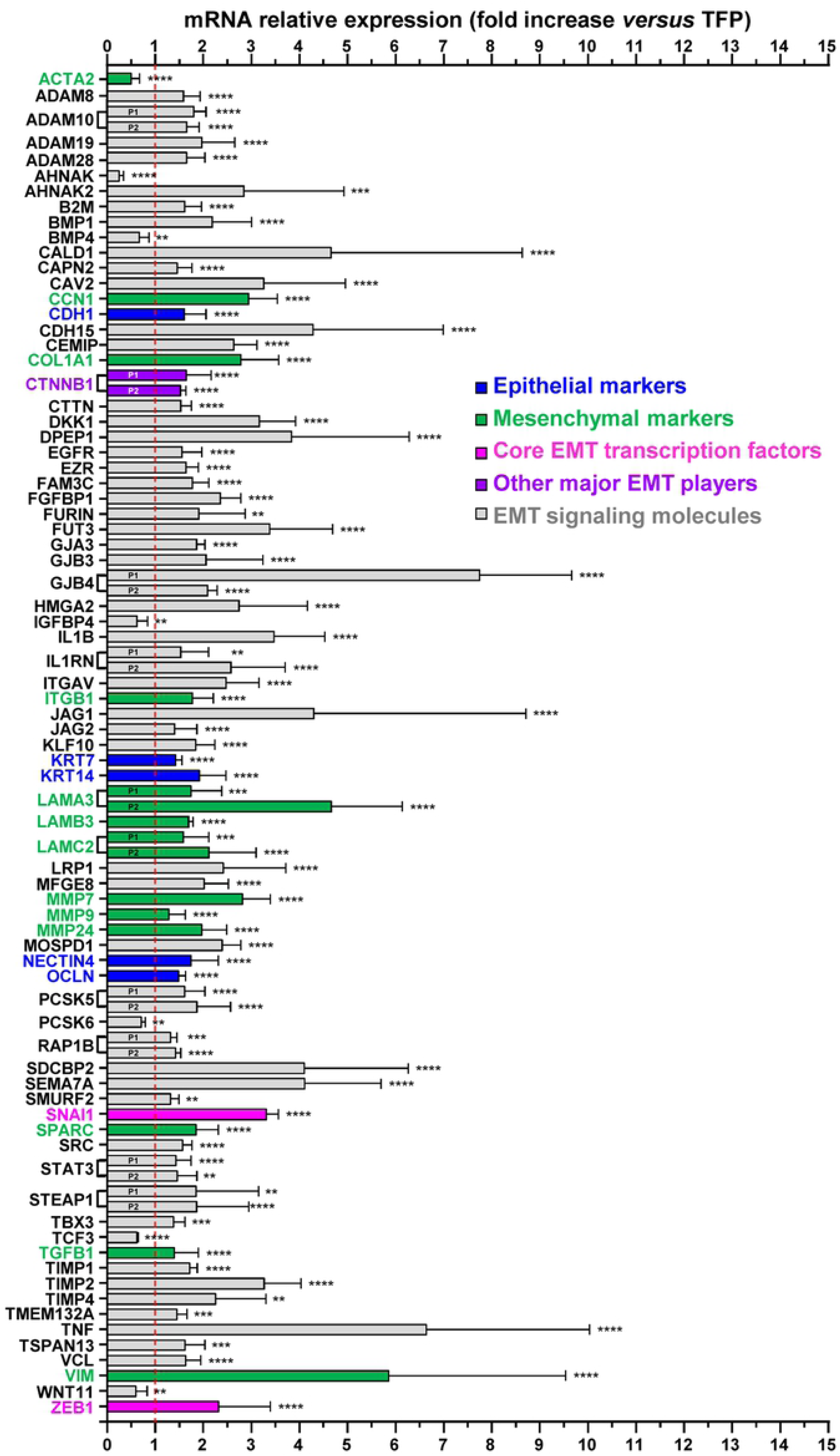
Expression profile of epithelial to mesenchymal transition-related genes differentially expressed in response to *Helicobacter hepaticus* CdtB in intestinal epithelial cells. The expression of genes was determined in HT-29 intestinal cells using the Human GE 4x44K v2 Microarray Kit (Agilent Technologies, Les Ulis, France) after a 72 h transduction with lentiviral particles expressing the CdtB subunit of *H. hepaticus* strain 3B1 *versus* the tdTomato fluorescent protein (TFP) as previously validated [13, 30]. The relative expression of genes in response to CdtB is reported as a fold change *versus* the value for cells cultured with lentiviral particles expressing the TFP. Results are presented as the mean ± SD of 4 replicates, as 4 independent transduction experiments were performed. The data presented for B2M, CAV2, CDH1, CTNNB1 (P1), EGFR, IL1B, ITGAV, ITGB1, JAG2, LRP1, MMP9, SEMA7A, SRC, and TGFB1 are the results of 40 replicates as 10 identical probes for each mRNA of these genes were included on the microarray Kit. The red discontinuous line shows the basal rate of GAPDH transcripts in cells expressing CdtB *versus* TFP. P1 and P2 represent the 2 probe names used for mRNA quantification. Details are presented in Table S2 (name and sequence of the probes, the corresponding gene name, the genbank accession number, the locus and the transcript variant). Raw data are presented in Table S1. Asterisks denote significant results. Gene expression differences were considered statistically significant if they showed ≥30% difference in mean expression levels between samples from CdtB *versus* TFP control condition, and if their associated p-value was <0.01. **p<0.01, ***p<0.001, ****p<0.0001. Abbreviations (from GeneCards, the human gene database): ACTA2, Actin Alpha 2, Smooth Muscle; AHNAK, AHNAK Nucleoprotein; AHNAK2, AHNAK Nucleoprotein 2; B2M, Beta- 2-Microglobulin; BMP1, Bone Morphogenetic Protein 1, BMP4, Bone Morphogenetic Protein 4; CALD1, Caldesmon 1; CAPN2, Calpain 2; CAV2, Caveolin 2; CCN1, Cellular Communication Network Factor 1; CDH1, Cadherin 1; CDH15, Cadherin 15; CEMIP/KIAA1199, Cell Migration Inducing Hyaluronidase 1; COL1A1, Collagen Type I Alpha 1 Chain; CTNNB1, Catenin Beta 1; CTTN, Cortactin; DKK1, Dickkopf WNT Signaling Pathway Inhibitor 1; DPEP1, Dipeptidase 1; EGFR, Epidermal Growth Factor Recept; EZR, Ezrin; FAM3C, FAM3 Metabolism Regulating Signaling Molecule C; FGFBP1, Fibroblast Growth Factor Binding Protein 1; FURIN, Furin, Paired Basic Amino Acid Cleaving Enzyme; FUT3, Fucosyltransferase 3 (Lewis Blood Group); GJA3, Gap Junction Protein Alpha 3; GJB3, Gap Junction Protein Beta 3; GJB4, Gap Junction Protein Beta 4; HMGA2, High Mobility Group AT-Hook 2; IGFBP4, Insulin Like Growth Factor Binding Protein 4; IL1B, Interleukin 1 Beta; IL1RN, Interleukin 1 Receptor Antagonist; ITGAV, Integrin Subunit Alpha V; ITGB1, Integrin Subunit Beta 1; JAG1, Jagged Canonical Notch Ligand 1; JAG2, Jagged Canonical Notch Ligand 2; KLF10, Kruppel Like Factor 10; KRT7, Keratin 7; KRT14, Keratin 14; LAMA3, Laminin Subunit Alpha 3 ; LAMB3, Laminin Subunit Beta 3; LAMC2, Laminin Subunit Gamma 2; LRP1, LDL Receptor Related Protein 1; MFGE8, Milk Fat Globule EGF And Factor V/VIII Domain Containing; MMP7, Matrix Metallopeptidase 7; MMP9, Matrix Metallopeptidase 9; MMP24, Matrix Metallopeptidase 24; MOSPD1, Motile Sperm Domain Containing 1; NECTIN 4, Nectin Cell Adhesion Molecule 4; OCLN, Occludin; PCSK5, Proprotein Convertase Subtilisin/Kexin Type 5; PCSK6, Proprotein Convertase Subtilisin/Kexin Type 6; RAP1B, Member Of RAS Oncogene Family; SDCBP2, Syndecan Binding Protein 2; SEMA7A, Semaphorin 7A (John Milton Hagen Blood Group); SMURF2, SMAD Specific E3 Ubiquitin Protein Ligase 2; SNAI1, Snail Family Transcriptional Repressor 1; SPARC, Secreted Protein Acidic And Cysteine Rich; SRC, SRC Proto-Oncogene, Non- Receptor Tyrosine Kinase; STAT3, Signal Transducer And Activator Of Transcription 3; STEAP1, STEAP Family Member 1; TBX3, T-Box Transcription Factor 3; TCF3, Transcription Factor 3; TGFB1, Transforming Growth Factor Beta 3; Transforming Growth Factor Beta 3; TIPM1, TIMP Metallopeptidase Inhibitor 1; TIPM2, TIMP Metallopeptidase Inhibitor 2; TIPM4, TIMP Metallopeptidase Inhibitor 4; TMEM132A, Transmembrane Protein 132A; TNF, Transmembrane Protein 132A; TSPAN13, Tetraspanin 13; VCL, Vinculin; VIM, Vimentin; WNT11, Wnt Family Member 11; ZEB1, Zinc Finger E-Box Binding Homeobox 1.

### CdtB induces SNAIL and ZEB1 nuclear accumulation

As microarray data have shown a CdtB-dependent increase in SNAI1 and ZEB1 transcripts, immunocytofluorescence analyses were performed to localize and quantify the corresponding proteins. We used stable transgenic cell lines allowing the conditional ectopic expression of *H. hepaticus* CdtB and its corresponding mutated CdtB (H_265_L) lacking catalytic activity to examine the effects specifically related to the CdtB enzymatic activity [26,29,30].

Compared to cells expressing H_265_L CdtB, those expressing CdtB showed an increase in SNAIL nuclear expression (2-, 1.5-, 5.1- and 1.5-fold increase for intestinal HT-29, Caco-2, SW480 and hepatic Hep3B, respectively, Figs. 3A,D, S1A and S2A and magnification in boxes). With regards to ZEB1, the protein is mainly localized in the nucleus and in the cytoplasm of cells expressing H_265_L CdtB, while ZEB1 expression was significantly increased in the nucleus upon CdtB expression (1.25-, 1.55-, 1.6- and 1.2-fold increase for HT-29, Caco- 2, SW480 and Hep3B, respectively, Figs. 3B,D, S1A and S2A). ZEB1 upregulation following CdtB expression was also confirmed using xenograft mouse models (Fig. 3C,D).

**Fig. 3.**
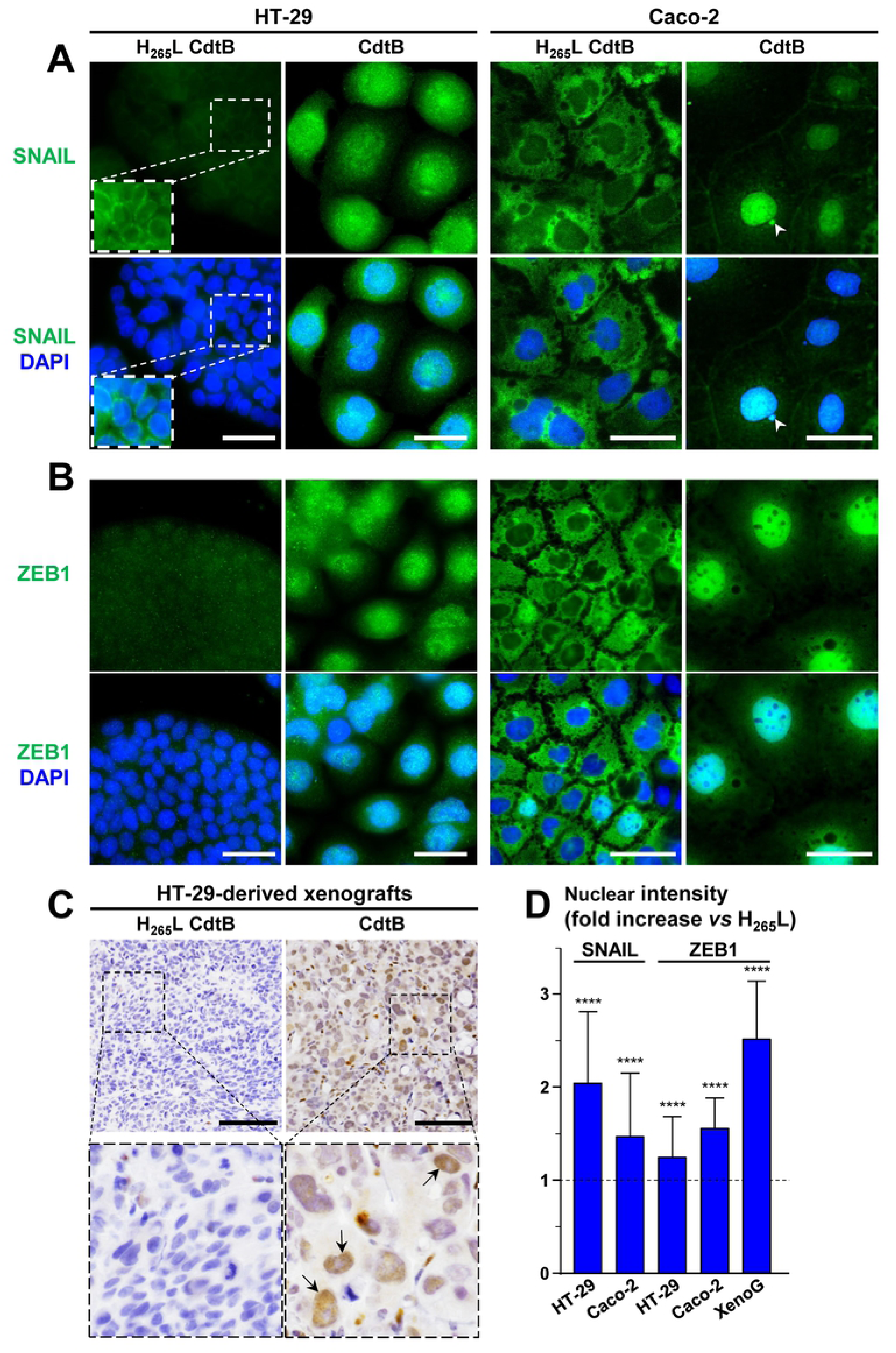
CdtB induces an increase in the nuclear expression of SNAIL and ZEB1 transcription factors. **(A)** and **(B)** Wide field images of intestinal epithelial cells HT-29 and Caco-2 cultured in the presence of doxycycline for 72 h to induce the expression of the CdtB of *H. hepaticus* strain 3B1 or its corresponding inactive form with the H_265_◊L mutation (H_265_L). Cells were processed for fluorescent staining with primary antibodies generated against SNAIL **(A)** or ZEB1**(B)** associated with fluorescent labeled-secondary antibodies (green) and DAPI to counterstain the nuclei (blue). A more intense color magnification is shown in boxes in (A). The white arrowheads indicates a micronucleus concentrating SNAIL. Scale bar, 40 µm. **(C)** Images of 3 μm-tissue sections of HT-29-CdtB- and HT-29-H_265_L-derived mice engrafted tumors immunostained for ZEB1 and counterstained with standard hematoxylin staining. HT29-transgenic cell lines were engrafted into immunodeficient mice as previously reported [26]. Magnifications of selected areas are shown in boxes. Black arrows indicate some nuclei concentrating ZEB1. Scale bar, 100 µm. **(D)** The nuclear intensity of SNAIL and ZEB1 transcription factors was quantified and is presented as fold change *versus* H_265_L. The results are presented as the mean ± SD of triplicates in one representative experiment out of three. The discontinuous line shows the basal rate in cells expressing the inactive form of the CdtB of *H. hepaticus* strain, H_265_L. ****p<0.0001 *versus* H_265_L. XenoG, xenograft.

Taken together, these data showed that the active CdtB subunit, promotes increased expression of EMT-TFs, SNAIL and ZEB1, as well as their increased nuclear translocation suggesting an increase in their pro-EMT activity.

### CdtB promotes the loss of epithelial cell junctions and individualization of cells

Epithelial cells are interconnected *via* epithelial cell junctions, including tight junctions, Adherens junctions, desmosomes and gap junctions. During EMT, cell–cell junctions are lost. We evaluated cell–cell junctions of epithelial cells by immunocytofluorescence. Analysis of cell junctions in the SW480 and Hep3B cell lines was inconclusive as these cells are poorly cohesive and their labeling appeared diffuse. Thus, this analysis was performed only in HT-29 and Caco-2 cells. Tight junction proteins are a hallmark of epithelial cells. The tight junction protein Zona Occludens protein 1 (ZO-1) is present at the plasma membrane in tightly cohesive HT-29 and Caco-2 cells expressing the inactive mutated H_265_L CdtB (Fig. 4A). Upon CdtB intoxication, ZO-1 disappeared from the junctions and appeared diffuse in the cytoplasm of HT-29 cells that appeared individualized. In CdtB intoxicated Caco-2 cells, ZO-1 completely disappeared over entire fields and the staining intensity showed a significant decrease (Fig. 4D), suggesting ZO-1 degradation. Cell-cell Adherens junctions are essential for the maintenance of epithelial homeostasis and can limit cell movement and proliferation. Immunofluorescent labeling of E-cadherin from Adherens junctions showed its presence at the plasma membrane in cohesive HT-29 and Caco-2 cells expressing the inactive H_265_L CdtB. In HT-29 cells expressing CdtB, a loss of E-cadherin in the junctions was observed and the protein was found relocated in the cytoplasm (Fig. 4B). A perinuclear localization of E-cadherin was also observed in some CdtB-intoxicated HT-29 and Hep3B cells (arrowheads in Figs. 4B and S2B). Analysis of E-cadherin expression in the mouse xenograft models showed an overall decrease in E- cadherin expression in some fields in response to CdtB (Fig S3A). Desmosomes junctions are intercellular adhesive junctions, which are anchorage points for intermediate filaments providing strong cell-cell adhesion. As observed for tight junctions and Adherens junctions, desmosomes were also disassembled in response to CdtB (Fig. 4C), leading to residual intercellular filaments of γ-Catenin (arrowheads in Fig. 4C). Quantification analyzes showed that CdtB significantly triggers a loss of cell-cell junction with the disassembly and removal of existing cell junctions (Fig. 4D).

**Fig. 4.**
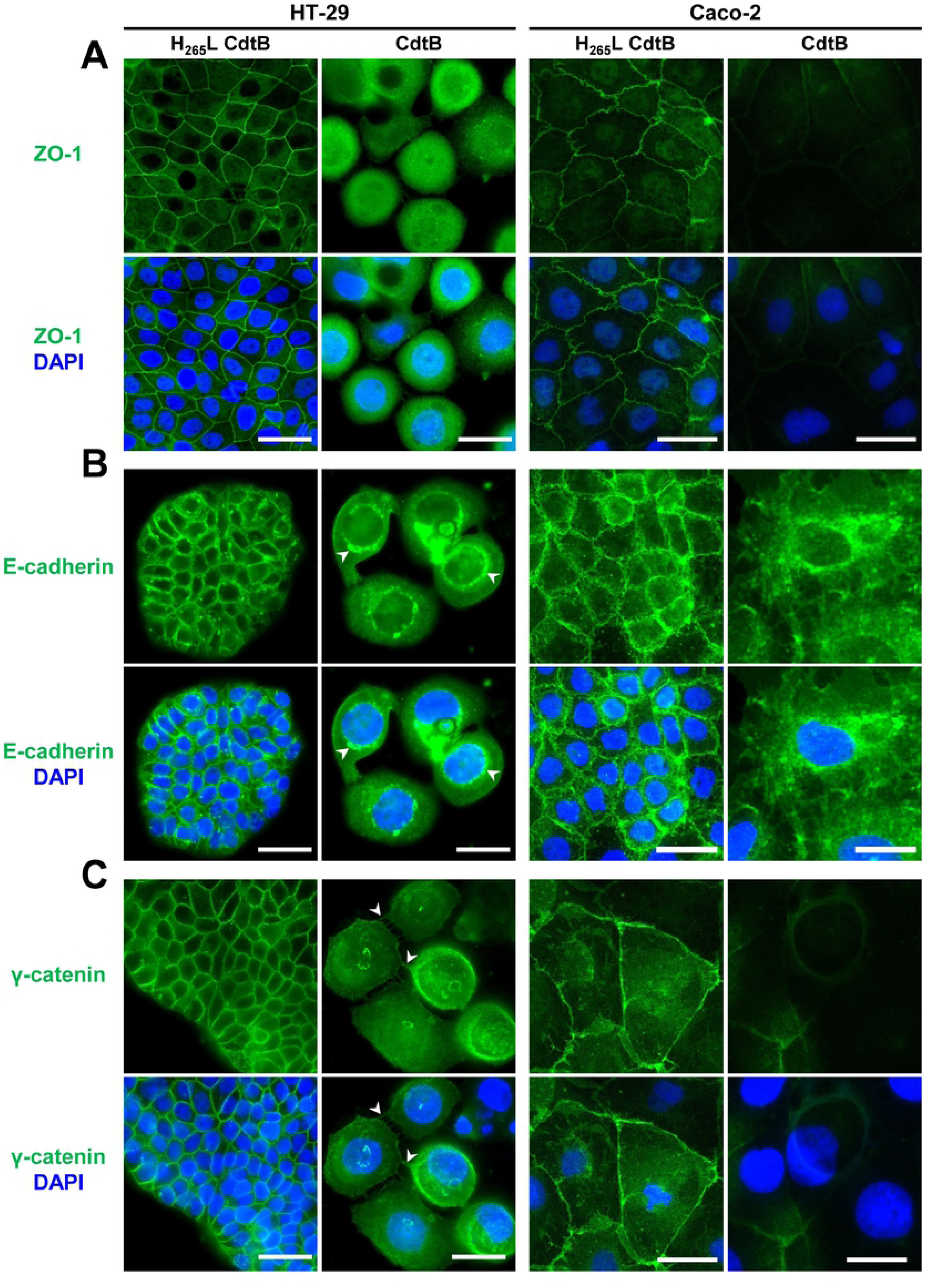
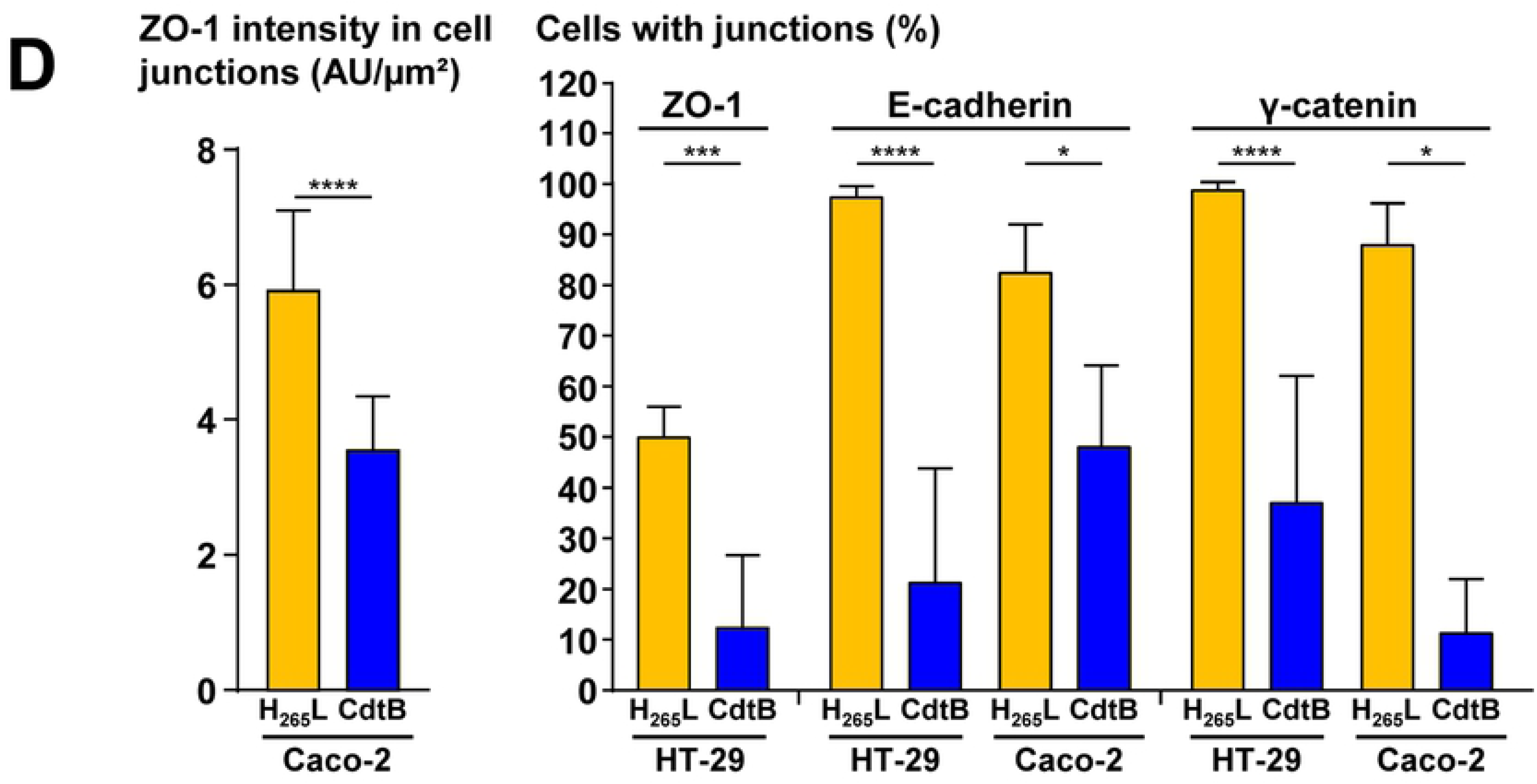
CdtB induces the loss of cell-cell junctions. **(A)** to **(C)** Wide field images of tight junctions (zonula occludens 1, ZO1), Adherens junctions (E-cadherin), and desmosomes (γ- catenin/plakoglobin), respectively, of HT-29 and Caco-2 intestinal epithelial cells cultured in the presence of doxycycline for 72 h to induce the expression of the CdtB of *H. hepaticus* strain 3B1 or its corresponding inactive form with the H_265_→L mutation (H_265_L). Cells were processed for fluorescent staining with primary antibodies generated against cell junctions associated with fluorescent labeled-secondary antibodies (green) and DAPI to counterstain the nuclei (blue). Scale bar, 40 µm. Arrowheads in (B) indicate the E-cadherin concentration and perinuclear relocalization in some CdtB-intoxicated HT-29 cells. Arrowheads in (C) indicate residual membrane spans containing γ-catenin. (**D**) Quantification of the loss of cell junctions. The percentage of cells displaying junction was counted manually, with the exception of ZO-1 labeling in Caco-2 cells. In those latest cells, ZO-1 quantification was performed by measuring the intensity of labeling within cell junctions and was expressed in arbitrary units per µm² (AU/µm²). A minimum of 500 cells was counted. The results are presented as the mean ± SD of triplicates in one representative experiment out of three. *p<0.05, ***p<0.001, ****p<0.0001 *versus* H_265_L.

During EMT, the loss of junctional proteins leads to individualization of cells which next display spindle-shaped morphology (also called mesenchymal-like phenotype). Some individualized and elongated CdtB cells were indeed observed at the periphery of HT-29 cell clusters (yellow and white arrowheads in Fig. 5A, respectively). This suggests that the CdtB-induced loss/delocalization of junctional proteins is accompanied by individualization of cells. Measurement of cell elongation in response to CdtB was challenging since CDT/CdtB-induced megalocytosis hindered this measurement. However, it was possible to observe and quantify the elongation of the morphology in SW480 cells. Indeed, these cells showed a significant cellular elongation as early as day 6 following the expression of CdtB (Fig. S1B).

**Fig. 5.**
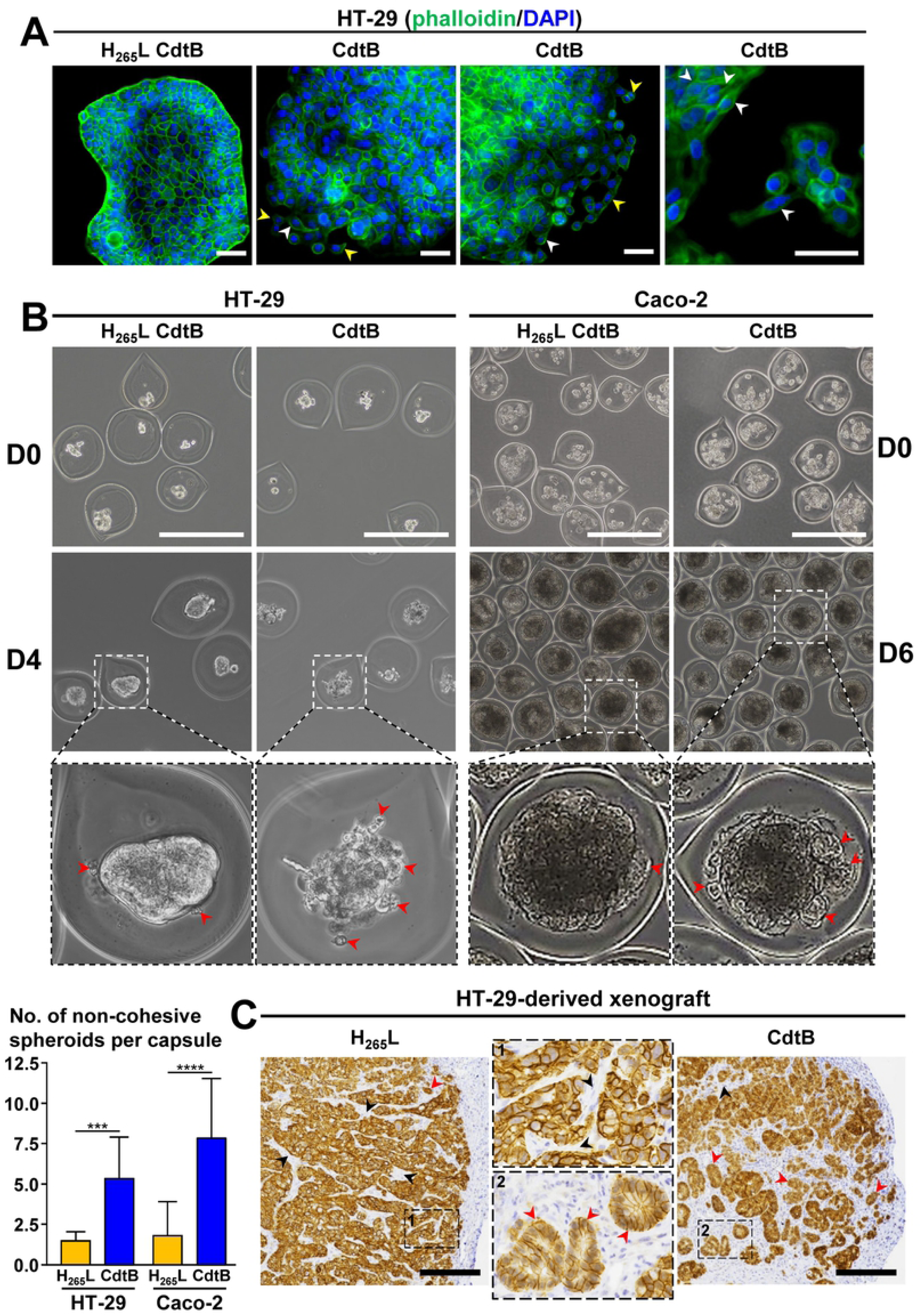
CdtB induces a loss of cellular cohesion and cell individualization. **(A)** Wide field images of intestinal epithelial cells HT-29 cultured in the presence of doxycycline for 72 h to induce the expression of the CdtB or H_265_L CdtB. Cells were stained with fluorescent-labeled phalloidin (green) to detect F-actin and DAPI to counterstain the nucleus. Scale bar, 50 µm. Yellow and white arrowheads indicate individualized and mesenchymal-like cells, respectively. **(B)** Analysis of the effects of CdtB on cell spheroids. Approximately 30 transgenic HT-29 and Caco-2 cells were encapsulated in alginate microcapsules and cultured during 24 and 48 hours, respectively. Then, doxycycline was added to induce the expression of CdtB or its corresponding inactive form with the H_265_→L mutation (H_265_L). For Caco-2 cells, doxycycline was added during 5 days. Then, new medium without doxycycline was added for 24 additional hours (day 6). The number of non-cohesive spheroid per capsule was counted manually (n=100 capsules) at day 4 and 6 for HT-29 and Caco-2, respectively. Data represent the mean ± SD of triplicates in 1 representative experiment out of 3. Scale bar, 400 µm. Red arrowheads indicate cells/spheroids individualizing inside the capsule. D, day. ***p<0.001, ****p<0.0001 *versus* H_265_L **(C)** Images of 3 μm-tissue sections of HT-29-CdtB- and HT-29-H_265_L-derived mice engrafted tumors immunostained for E-cadherin, and counterstained with standard hematoxylin staining (blue). HT-29-transgenic cell lines were engrafted into immunodeficient mice as previously reported [26]. Magnifications of selected areas are shown in boxes. Black and red arrowheads indicate mice stromal cell infiltrates and individualized cell clusters at the periphery of HT-29- CdtB-derived tumor. Scale bars, 250 μm.

To study intercellular interactions in a three-dimensional environment mimicking the characteristics of a tissue, we used cells encapsulated in alginate microcapsules, having the property of being permeable to different molecules (Fig. 5B). Thus, after encapsulation of approximately 30 cells per capsule, transgene expression was induced by addition of doxycycline. After four days of transgene induction, the HT-29 derived-spheroid expressing H_265_L CdtB grew up with a well-delineated spheroid in the capsule than those expressing CdtB. In the capsules in which the cells express CdtB, spheroids were split into several smaller individualized spheroids. With regards to Caco-2 derived-spheroid, those expressing CdtB have budded and are sometimes individualized. Quantification of the number of spheroids present within each alginate capsule showed a significant increase in the number of spheroids upon CdtB expression, suggesting that, as in the two-dimensional cell culture, CdtB causes cell individualization by a loss of cellular cohesion. Less cohesive and smaller individualized cell clusters were also observed at the periphery of HT-29-CdtB-derived tumor (Fig. 5C).

### CdtB increases Fibronectin, α5β1 integrin and matrix metalloproteinases expression and activity

As cells detach from the epithelium and extend lamellipodia from their basal surface, they next breach the basal lamina barrier, a layer of ECM secreted by epithelial cells. Fibronectin is an ECM glycoprotein and a mesenchymal marker whose primary receptor is α5β1 integrin. Altered expression of Fibronectin, α5β1 integrin and MMPs is a hallmark of invasive tumor cells.

Fibronectin transcript and protein were not detected at the basal level in HT-29 cell line (S1 Table and Fig. 6A) while increased fibronectin expression was shown in Caco-2 cells upon CdtB expression. α5 and β1 subunits of α5β1 integrin were also found upregulated in HT-29 and Caco-2 cells expressing CdtB (Fig. 6A). MMPs are secreted zinc-dependent endoproteases that degrade the protein components of the ECM. During EMT, the expression of several proteases increases, including MMPs. The expression of MMPs and their proteolytic activity were analyzed.

**Fig. 6.**
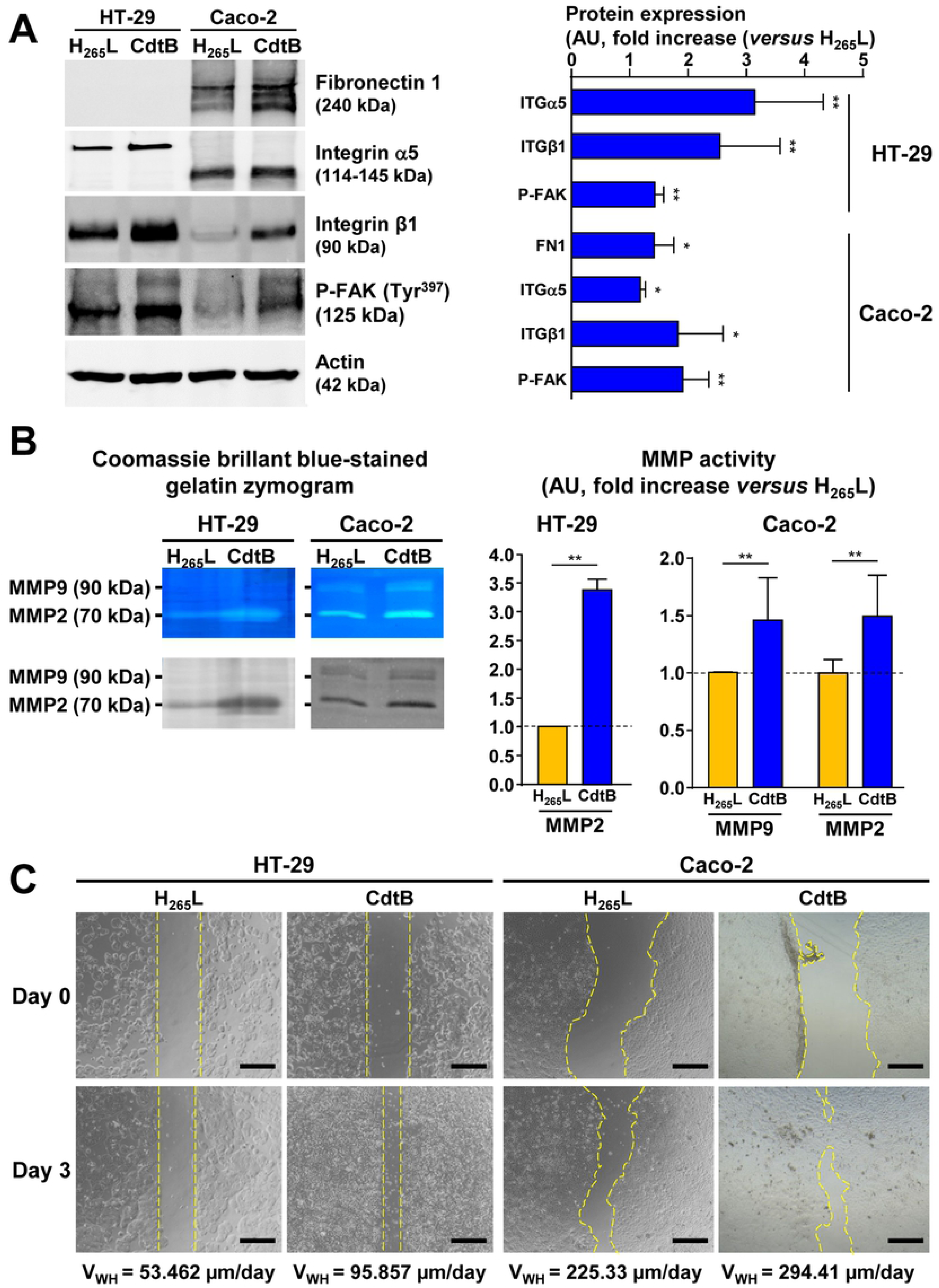
CdtB increases matrix metalloproteinases activity and cellular motility of naive cells. **(A)** Western blot analysis of the protein expression level of Fibronectin 1, Integrin α5, Integrin β1 and Phospho-FAK (Tyr397)following 3-days of CdtB and H_265_L expression induction in transgenic HT-29 and Caco-2 cells. Fibronectin was not detected in HT-29 cells. Data represent the mean ± SD from 3 independent experiments. **(B)** The activity of the MMP was determined by gelatin degradation zymography assay. HT-29 and Caco-2 cells were cultured in the presence of doxycycline for 72 h to induce the expression of the CdtB of *H. hepaticus* strain 3B1 or its corresponding inactive form with the H_265_→L mutation (H_265_L). Samples (cells and supernatant) were collected 3 days after the end of transgene induction, i.e., on day 6 after doxycycline induction, with the culture medium being replaced by 1 mL of medium without fetal calf serum (FCS) on the fifth day after induction, as FCS has a strong metalloprotease activity. Proteins were subjected to gelatin zymography to quantify MMP9 and MMP2 activity. As the analysis of supernatants was not conclusive, only MMP activity of the cell lysates is presented. The relative MMP activity in cells was determined by densitometric analysis of the transparent gelatin degradation bands/strips in Coomassie Brilliant Blue-stained zymograms, using Fiji (v1.53j) software. MMP activity is presented as fold change *versus* H_265_L. The results are presented as the mean ± SD of triplicates. The discontinuous line shows the basal rate of MMP activity in cells expressing the inactive form of the CdtB of *H. hepaticus* (H_265_L). **p<0.01, *versus* H_265_L. **(C)** Analysis of the migration induced by the secretome of cells expressing CdtB and its inactive form H_265_L. Confluent naive HT-29 and Caco-2 cells were scratched, washed to remove cellular debris and subsequently treated for a duration of 3 days with freshly collected supernatant of transgenic HT-29 and Caco-2 cells expressing CdtB and H_265_L CdtB (3-days doxycycline treatment). Images were acquired daily under a phase-contrast microscope to be further analyzed using Fiji (v1.53j) to determine the distance between the wound borders. A closing curve (distance/day) was constructed for each experimental conditions (n=6) and the slope of the fitting trend curve calculated to determine the wound healing velocity. On the phase contrast micrographs showing the wound-healing experiment in HT-29 and Caco-2 cells, the dotted lines define the borders of the wound. Scale bar, 500 µm. Abbreviations: AU, arbitrary units; D, day; ITGα5, integrin α5 subunit; ITGβ1, integrin β1 subunit; MMP, matrix metalloproteinase; P-FAK, Phospho-FAK (Tyr^397^) ; V_WH_, wound healing velocity.

In HT-29 cells, microarray data showed a CdtB-dependent upregulation of some MMP transcripts, i.e. *MMP7*, *MMP9* and *MMP24* (*MT5-MMP*) (Fig. 2) while *MMP2* was not detected at the basal level using the probe used for microarray analysis. As different *MMP2* alleles were reported, RT-qPCR with primers designed on a conserved region of MMP2 was performed and showed the upregulation of *MMP2* in HT-29 (Fig. S3B). CdtB also increased the transcription of *FURIN*, *PCSK5* and *PCSK6* pro-convertases, involved in the activation of MMPs *via* proteolytic cleavage of their pro-MMP precursors. Additionally, the mRNA level of Tissue Inhibitor of Metalloproteinases *TIMP1*, *TIMP2* and *TIMP4*, increased also upon CdtB expression. In Caco-2 cells, RT-qPCR showed the upregulation of *MMP2*, *MMP7* and *MMP9* following CdtB expression in (Fig. S3B), in contrast to and *MMP24* (Fig. S3B). Gelatin degradation assay revealed a significant increase in MMP2 proteolytic activity in both HT-29 and Caco-2 cells and MMP9 in Caco-2 cells, detected by gelatin zymography (Fig. 6B).

### The secretome of CdtB-intoxicated cells induces an increase in cellular motility

Wound-healing assay was next performed to study directional cell migration *in vitro*. Unfortunately, the expression of CdtB induced a cell cycle arrest and significant cell death [31, 32], leading to non-conclusive wound-healing assays. As CdtB-expressing cells secrete several pro-EMT factors (BMP1, IL1β, TGFβ1 and TNF, Fig. 2), the impact of the supernatant from CdtB-expressing cells on migration properties was evaluated on naive in HT-29 and Caco- 2 cells. As shown in Fig. 6C, accelerated wound healing was measured in the presence of the supernatant from cells expressing ‘active CdtB’ than with that from cells expressing ‘mutated H_265_L CdtB’. This suggests that the secretome induced by CdtB stimulates the closure of the wound, thus increases cellular motility.

## Discussion

Ever since the demonstration of the role of *Helicobacter pylori* infection in the development of gastric adenocarcinoma in humans *via* its CagA oncoprotein [33], other bacteria-producing toxin have been reported [34] to contribute to cancer hallmarks. This study investigated the main steps characterizing the EMT process following the intoxication of epithelial cells with the cytolethal distending toxin of *H. hepaticus*. The main findings of this study are summarized as follows: 1) *H. hepaticus* infection is associated with increased Vimentin and nuclear TWIST expression along with decreased E-cadherin expression in the liver of infected mice; 2) CdtB subunit of *H. hepaticus* CDT elicits EMT process in 2D epithelial cell lines; 3) CdtB-induced EMT features have been also observed in 3D models: xenografts and cells spheroids encapsulated in alginate microcapsules. Overall, most of EMT features were observed upon CdtB expression in the different models implemented in this study. Epithelial cells that undergo EMT lose their epithelial phenotype to acquire mesenchymal characteristics and become migratory and invasive. The loss of epithelial markers and gain of mesenchymal markers was not complete upon CdtB intoxication. Indeed, microarray microarray analysis did not evidence CdtB-induced downregulation of epithelial transcripts but showed a CdtB-upregulation of numerous EMT signaling transcripts, an increase in mesenchymal transcripts, as well as those from β-catenin and EMT TTs, SNAIL and ZEB1, . Thus, mixed epithelial and mesenchymal genes are expressed upon *H. hepaticus* infection and CdtB expression. Similar results were observed at protein level in the various models implemented in this study. These results are not surprising since 1) tumor cells are heterogeneous and in are in multiple transitional states and 2) EMT is not complete in cancer cell lines [35]. Indeed, cancer cells undergoing EMT are frequently found in a hybrid state known as partial EMT where epithelial cells express both epithelial markers and mesenchymal ones [36]. At protein level, most of major EMT transcriptional regulators, TWIST, SNAIL and ZEB1, were found overexpressed. ZEB1 is required for DNA repair and the clearance of DNA breaks and ensures the maintenance of genomic stability over the course of tumorigenesis [37]. Thus, the DNA damage induced by CdtB would trigger ZEB1 expression to prevent genomic instability. Following CdtB intoxication, other EMT markers were upregulated (Vimentin, Fibronectin, α5β1 Integrin) and FAK was phosphorylated. Activation of β1 integrin leading to enhanced cell spreading on fibronectin upon CDT effect was recently reported [38]. It was previously reported that 1) mesenchymal cells express Vimentin intermediate filaments and attach to ECM *via* integrin-containing focal adhesions [24] and 2) Fibronectin promotes adhesion and migration through integrin α5β1-mediated phosphorylation of FAK [39]. By upregulating and activating these proteins expression and FAK phosphorylation, CdtB might promote attachment to ECM and migration. Indeed, at the onset of migration, cells often exhibit increased Vimentin expression which protects cells against nuclear rupture and DNA damage during migration [40] and regulates nuclear shape, perinuclear stiffness, cell motility, as well as the ability to resist large cellular deformations [40]. Accordingly, the nuclease activity of CdtB would allow the recruitment of Vimentin to protect CdtB-intoxicated cells from DNA damage and control the size of their nucleus and to go through cell megalocytosis, enabling cell survival following the CdtB-induced DNA damage.

Depending on the model, CdtB induced the disassembly and removal of junctional proteins with their subsequent relocation (ZO-1 from tight junctions, E-cadherin from Adherens junctions, γ-Catenin/Plakoglobin from desmosomes). Fields of decreased expression of junctional proteins were observed both *in vivo* upon *H. hepaticus* infection and *in vitro* upon CdtB expression, despite an unexpected CdtB-induced increase in CDH1 mRNA expression. Substantial alteration of the cytosolic distribution of E-Cadherin was previously observed in human gingival explants following the exposure to the CDT of *Aggregatibacter actinomycetemcomitans* [41]. This CDT-induced Adherens junctions remodeling was associated to a pronounced increase in the expression and cytosolic distribution of E-cadherin [41]. The decrease of E-cadherin during EMT is not systematic. Indeed, cells who underwent EMT following the intoxication with *Escherichia coli* protein toxin cytotoxic necrotizing factor 1 (CNF1) did not show reduced expression of E-cadherin, but the latter was relocated from membrane to the cell body [42]. The loss of E-cadherin in Adherens junctions following CdtB expression was accompanied with its relocation in the cytoplasm as well as at perinuclear region (Fig. 4B and S2B). Accumulation of E-cadherin bound to β-catenin at the perinuclear endocytic recycling compartment upon dissociation of Adherens junctions was reported and both proteins can be translocated into the nucleus upon Wnt pathway activation [43]. Thus, perinuclear E-cadherin relocation following CdtB expression suggests Wnt pathway activation. In addition to its cellular internalization, E-Cadherin is also cleaved by MMPs and ADAM proteins, a proteolysis known to facilitate E-Cadherin ectodomain shedding to further destabilize junctions [44]. Tight junctions can be also internalized *via* Caveolin-mediated endocytosis [45]. Thus, the CdtB-increase of MMPs, ADAMs and CAV2 mRNA level would reflect increased degradation of Adherens junctions and tight junctions proteins. The loss of cellular junctions is concomitant with cell individualization.

Another feature of EMT is the remodeling of actin cytoskeleton and reorganization of stress fibers. These phenotypes have already been investigated and reported that CDT induce a profound remodeling of the actin cytoskeleton with the formation of stress fibers [20]. We previously reported that CDT/CdtB induces an atypical delocalization of vinculin from focal adhesions to the perinuclear region, formation of cortical actin-rich large lamellipodia with an upregulation of cortactin protein [21, 46]. Reorganization of the actin cytoskeleton and focal adhesion formation are hallmarks of EMT, resulting in increased cell movement. Cortactin gene and protein were also overexpressed by CdtB (Figs. 2 and S3D) which triggers the formation of large lamellipodia (Fig. S3C) and fibroblast-/mesenchymal-like morphology (Fig. 5A and S1B). These results suggest that CdtB induces the reorganization of the actin cytoskeleton to acquire more spindle-like morphology, a phenotype known to allow polarized assembly of the cytoskeleton into protrusive and invasive structures that help cells to move through ECM [47]. In this context, CdtB-induced lamellipodia formation would be enhanced by the expression of cortactin whose expression facilitates lamellipodia formation and contributes to the maintenance of migration persistence *via* stabilisation of the actin network [48].

A variety of factors in microenvironment can lead to EMT. Here, we also showed CdtB increase the expression of pro-EMT cytokines and the secretome of CdtB-expressing cells increases cellular motility of naive cells, emphasizing the important effect of CdtB-induced secretome on cellular microenvironment. Bacterial genotoxins enables cells to release pro- inflammatory molecules and harbor a senescence-associated secretory phenotype (SASP) [26, 49] known to modify the cellular microenvironment [50, 51]. Knowing that inflammation and/or SASP-induced chemokines are key inducers of EMT changes in neighboring cell populations [52], chronic exposure to CDT/CdtB might promote an EMT process in neighboring cells in a CdtB inflammatory environment. In line with that, the increase in Vimentin expression observed in sinusoidal endothelial cells in the liver of *H. hepaticus*- infected mice (Fig. 1C) could be driven by the secretome of infected hepatocytes.

Bacterial genotoxins, CDT and colibactin, cause genome instability whose persistence lead to tumorigenic phenotypes. The molecular mechanisms enabling malignant progression involve EMT-dependent barrier alterations and tumor-promoting inflammation. Several bacterial toxin elicit EMT process, i.e. CagA cytotoxin of *H. pylori* [53], α-toxin/hemolysin A of *S. aureus* [54], CNF1 of *E. coli* [42] and *Bacillus fragilis* toxin [55]. With regards to bacterial genotoxins, previous data have shown that bacterial genotoxins induce certain phenotypes reminiscent of EMT. Collectively, our *in vivo* and *in vitro* findings present evidence consistent with a CDT/CdtB-induced EMT process. However, the signaling pathways involved in this process remain to be defined.

## Materials and methods

Antibodies and compounds used for immunohistochemistry, immunocytochemistry and western blot analyzes are presented in Table 1.

**Table 1.** Antibodies and compounds

| **Antibodies and compounds** | **Origin, clonality** | **clone** | **Reference, supplier, city, country** | **Dilution, pH, incubation** |
| --- | --- | --- | --- | --- |
| **Histology** |  |  |  |  |
| anti-E-Cadherin | Mouse, monoclonal | 36/E-Cadherin | #610181, BD Biosciences, San Jose, CA | dilution 1/200 pH9 1h |
| anti-TWIST | Rabbit, polyclonal | H-81 | #sc-15393, Santa Cruz Biotechnology, Heidelberg, Germany | 1/50 pH6 2h |
| anti-Vimentin | Mouse, monoclonal | V9 | #M0725, Agilent Dako, Santa Clara, CA | dilution 1/250 pH9 1h |
| anti-ZEB1 | Rabbit, polyclonal | / | #A301-922A, Bethyl Laboratories, Inc/Fortis Life Sciences, Montgomery, TX | dilution 1/100 pH9 2h |
| **Immunocytochemistry** |  |  |  |  |
| anti-Cortactin (p80/85) | Mouse, monoclonal | 4F11 | #05-180-I, Sigma-Aldrich | 1/100 |
| anti-E-Cadherin | Mouse, monoclonal | 36/E-Cadherin | #610181, BD Biosciences | 1/100 |
| anti-SNAI1 | Rabbit, polyclonal | H-130 | #sc-28199, Santa Cruz Biotechnology | 1/100 |
| anti-Vimentin | Mouse, monoclonal | V9 | #M0725, Agilent Dako | 1/100 |
| anti-ZEB1 | Rabbit, polyclonal | / | #A301-922A, Bethyl Laboratories, Inc/Fortis Life Sciences, Montgomery, TX | 1/100 |
| anti-ZO-1 | Rabbit, polyclonal | / | #61-7300, Invitrogen, Cergy Pontoise, France | 1/100 |
| anti-γ-Catenin/Plakoglobin | Rabbit, monoclonal | D9M1Q | 75550, Cell Signaling Technology distributed by Ozyme, Saint-Cyr-L'École, France | 1/100 |
| Alexa Fluor® 594-dye secondary antibody | Donkey, polyclonal | / | R37119, Molecular Probes, Invitrogen | 1/200 |
| Alexa Fluor® 488-dye secondary antibody | Donkey, polyclonal | / | R37118, Molecular Probes, Invitrogen | 1/200 |
| Phalloidin Alexa Fluor® 647 | / | / | A22287 Molecular Probes, Invitrogen | 1/500 |
| 4',6'-diamidino-2-phenylindol (DAPI) | / | / | D1306 Molecular Probes, Invitrogen | 1/1000 |
| **Western blot** |  |  |  |  |
| anti-fibronectin | Mouse monoclonal | 10 | #610077, BD Biosciences | 1/1,000 |
| anti-integrin alpha 5 | Rabbit, polyclonal | / | #GTX130705, GeneTex distributed by EUROMEDEX,<br>Souffelweyersheim, France | 1/2,000 |
| anti-integrin beta 1 | Rabbit, monoclonal | EP1041Y | #ab52971, abcam, Paris, France) (dilution | 1/1,000 |
| phospho-FAK (Tyr<sup>397</sup>) | Rabbit, monoclonal | D20B1 | #8556, Cell Signaling Technology | 1/1,000 |
| anti-β-actin | Mouse monoclonal | AC-74 | #A5316, Sigma Aldrich) (dilution | 1/5,000 |

- 1 For histology, anti-TWIST and anti-ZEB1 antibodies were used in combination with ImmPRESS® HRP Horse Anti-Rabbit IgG Polymer - 2 Detection Kit, Peroxidase (#MP-7401, Vector Laboratories, Burlingame, CA); anti-E-Cadherin was used in combination with ImmPRESS® HRP - 3 Horse Anti-Mouse IgG Polymer Detection Kit, Peroxidase (#MP-7402, Vector Laboratories); and anti-Vimentin was used in combination clone - 4 V9 with VECTASTAIN® Elite ABC-HRP Kit, Peroxidase, R.T.U. (Universal) (#PK-7200, Vector Laboratories).

### Ethics statement

Animal material provided from previous studies was approved by the Ethics Committee for Animal Care and Experimentation CEEA 50 in Bordeaux (Comité d’Ethique en matière d’Expérimentation Animale agréé par le ministre chargé de la Recherche, “dossier no. Dir13126BV2”, “saisines” no. 4808-CA-I [25] and 13126B [26], Bordeaux, France), according to treaty no. 123 of the European Convention for the Protection of Vertebrate Animals. Animal experiments were performed in an A2 animal facility (security level 2) by trained authorized personnel only.

### Transcriptomics

The global expression of human genes following the lentiviral ectopic expression of the CdtB subunit of *H. hepaticus versus* the tdTomato fluorescent protein (72 h transduction) was quantified in epithelial intestinal HT-29 cells using the Human GE 4x44K v2 Microarray Kit (Agilent Technologies) as we previously detailed [13, 30].

### Cell lines and culture conditions

The human HT29 (DSMZ collection no. ACC 299), Caco- 2 cells (DSMZ collection no. ACC 169) and SW480 (ATCC collection no. CCL-228) cell lines were derived from a colon adenocarcinoma, and the human epithelial cell line Hep3B (ATCC collection no. HB-8064) from a hepatocellular carcinoma. Their culture conditions are shown in [21, 29]. The corresponding transgenic CdtB- and H265L-cell lines were established as previously reported [21] and grown in their respective culture medium supplemented with 10% heat-inactivated fetal calf serum (Invitrogen) at 37°C in a 5% CO_2_ humidified atmosphere. When required, the transgene expression was induced in the cells from the tetracycline-inducible promoter by addition of doxycycline (200 ng/ml, #D9891, Sigma Aldrich Saint-Quentin Fallavier, France) to the culture medium.

### Immunodetection, image analysis and protein quantification

Cell cultures grown on glass coverslips in 24-well plates and fluorescently labeled were mounted on microscope slides with DAPI Fluoromount-G (Clinisciences SA, Montrouge, France). Dual- and triple-color imaging with Fluor 594-labeled phalloidin (red), Alexa Fluor 488-labeled (green) secondary antibodies, and 4’,6’-diamidino-2-phenylindole (DAPI, blue) was obtained using selective LED excitation at 594, 488 and 358 nm, respectively. Traditional widefield fluorescence imaging was performed using Eclipse 50i epi-fluorescence microscope (Nikon, Champigny sur Marne, France) equipped with the Nis Element acquisition software and a 640 (numerical aperture, 1.3) oil immersion objective (40X). The acquired images were calibrated according to the microscope software manufacturer. For immunohistochemistry, 3 μm-tissue sections of mice livers [25] and xenograft-derived tumors [26] were prepared from formalin-fixed paraffin-embedded tissues and submitted to standard hematoxylin staining and immunostaining using horseradish peroxidase coupled antibodies and 3,3’-diaminobenzidine (DAB).

For quantification of nuclear EMT protein markers, nuclear surface was delineated by isolating the nuclear counterstaining (hematoxylin or DAPI staining) for each nucleus by using the ‘Threshold’ function of Fiji (v1.53j) software [56]. Then, protein quantification in nucleoplasm was performed by measuring the sum of pixel intensity with the “Raw integrated density” measure function of Fiji [56] using the delineation of hematoxylin staining or DAPI fluorescence for each nucleus. A minimum of 1,000 nuclei was measured.

For the percentage of stained surface measured in immunohistochemistry, the hematoxylin and DAB staining were separated using the “Color Deconvolution” function of Fiji. The DAB stained surface was then measured using the “Threshold” function and a ratio made with the whole tissue surface.

### Cell encapsulation in alginate microcapsules

Cell encapsulation was performed as previously described by our team using the microfluidic technique to produce 3D cell-based assays with minor modifications [57]. First, the microfluidic device was 3D printed and a glass capillary (∼150 µm diameter) was glued to the exit of the device. Then three solutions loaded in syringes mounted to pumps were injected into the coaxial cones of the device. The outermost cone contained the alginate solution; the intermediate cone contained a 300 mM sorbitol solution, and the innermost cone contained the cells in sorbitol or Matrigel/sorbitol solution. Composite droplets exiting the nozzle fell in a 100 mM CaCl_2_ bath for gelling on. Capsules were washed with cell culture medium and subsequently cultured in the medium recommended for the cell line encapsulated.

### Zymography assay

Gelatinolytic activity of MMP was examined as previously described [18, 19]. Briefly, 8% polyacrylamide gels, co-polymerized with gelatin (2 mg/ml), were loaded with 50 μg of protein collected 3 days after the end of transgene induction, i.e., on day 6 after induction. After electrophoresis, gels were washed 4 times with renaturation buffer (containing 1% Triton X-100) to remove sodium dodecyl sulfate, and then incubated at 37°C for 48 h in an incubation buffer containing the cofactors necessary for the activation of gelatin degradation proteins (2.5% Triton X100, 50 mM Tris HCl, 10 mM CaCl2, and 0.2 M NaCl). Then, the gels were stained with 0.05% Coomassie Brilliant Blue, and gelatinolytic activities were detected as transparent bands against the dark-blue background. Quantification of proteolysis (band intensity) was performed using Fiji (v1.53j) software and expressed in arbitrary units.

### Two-steps wound-healing assay

First, transgenic CdtB and H_265_L HT-29 and Caco-2 cells were seeded in 24-well plate at a density of 4x10^4^ cells/well for 24 h. The cells were then treated with doxycycline for a duration of 4 days to induce the expression of the CdtB and inactive H_265_L CdtB; the supernatants were collected and the cells were discarded. Second and in a separate experiment initiated on the third day of the first experiment, naive (non-transgenic) HT-29 and Caco-2 cells were seeded in 24-well plate at a density of 5x10^5^ and 2x10^5^ cells/well, respectively, until confluency (24 h). Then, the cell monolayers were scraped in a straight line to create a “scratch” with a p10 pipet tip. Cells were washed 3 times with 1 ml of culture medium to remove the cellular debris and 1 ml of the supernatant freshly collected from transgenic CdtB- and H_265_L-first culture was added and incubation was continued for three days. For each sample (CdtB- and H_265_L), images were acquired daily at the same field of view using a phase-contrast microscope to be further analyzed quantitatively with Fiji (v1.53j) software to analyze the effect on the motility induced by the secretome of cells expressing CdtB and inactive CdtB and determine the average wound size. The closure curve was then generated on Excel and the slope of the fitting trend curve was used to determine the closure speed.

## Acknowledgments

1) This research was funded in part by the ‘Ligue Contre le Cancer Gironde’. We are indebted to Wencan He for technical assistance in preliminary experiments. The authors thank Laetitia Andrique for technical assistance in alginate cell encapsulation. The 3D capsules were obtained from the VoxCell Facility, a service of the CNRS-INSERM and Bordeaux University - UMS TBMCore 3427. Lornella Seeneevassen is the recipient of a predoctoral fellowship from the French Ministry of Education, Research and Scientific Innovation. Ruxue Jia is the recipient of a predoctoral fellowship from China Scholarship Council.

2) Part of this work was presented as oral presentation at the ETOX european workshop on bacterial protein toxins, 2021.

## Conflict of interest

The authors disclose no conflicts.

## Supporting information

**S1 Fig. Effects of CdtB on EMT markers in SW480 cells.** Intestinal epithelial SW480 cells were cultured in the presence of doxycycline for 72 h to induce the expression of the CdtB of *H. hepaticus* strain 3B1 or its corresponding inactive form with the H_265_→L mutation (H_265_L). **(A)** Three days after doxycycline induction, cells were processed for fluorescent staining with primary antibodies generated against EMT TFs, i.e. SNAIL and ZEB1, associated with fluorescent labeled-secondary antibodies (green) and DAPI to counterstain the nuclei (blue). The nuclear intensity of SNAIL and ZEB1 TFs was quantified and is presented as fold change *versus* H_265_L. The discontinuous line shows the basal rate in cells expressing H_265_L. The results are presented as the mean ± SD of triplicates in one representative experiment out of three. **(B)** Three days after doxycycline induction, incubation was continued for 3 to 5 days corresponding to day 6 to day 8 with respect to the first day of doxycycline induction. Cells were then stained daily with fluorescent-labeled phalloidin (pink) and DAPI (blue) to detect F-actin and the nucleus, respectively. Cell elongation (length/width ratio measurement) was performed using Fiji software (n=100). Images are from a representative manipulation. White arrowheads indicate mesenchymal-like cells. Scale bar, 40 µm. *p<0.05, **p<0.01, ***p<0.001 *versus* H_265_L.

**S2 Fig. Effects of CdtB on EMT markers in Hep3B cells.** Hepatic epithelial cells Hep3B were cultured in the presence of doxycycline for 72 h to induce the expression of the CdtB of *H. hepaticus* strain 3B1 or its corresponding inactive form with the H_265_→L mutation (H_265_L). Cells were immunostained with fluorescent antibodies (green) targeting EMT TFs **(A)**, i.e. SNAIL and ZEB1, and E-Cadherin **(B)**, and DAPI to counterstain the nucleus (blue). The nuclear intensity of SNAIL and ZEB1 TFs was quantified and is presented as fold change *versus* H_265_L. The discontinuous line shows the basal rate in cells expressing H_265_L. The results are presented as the mean ± SD of triplicates in one representative experiment out of three. The yellow and white arrowheads in **(B)** indicate residual membrane spans containing E-Cadherin and perinuclear localization of E-cadherin, respectively. Scale bars, 10 µm. ***p<0.001 *versus* H_265_L. **(C)** Images of 3 μm-tissue sections of Hep3B-CdtB- and Hep3B-H_265_L-derived mice engrafted tumors immunostained for E-cadherin and Vimentin (brown), and counterstained with standard hematoxylin staining (blue). Hep3B-transgenic cell lines were engrafted into immunodeficient mice as previously reported [26]. Magnifications of selected areas are shown in boxes. Black arrows indicate enlarged cells presenting less junctional E-cadherin. Red arrows indicate Vimentin invaginated in the nucleus of giant cells, corresponding to nucleoplasmic reticulum [29]. Scale bar, 100 µm.

**S3 Fig. Other CdtB effects on EMT markers in intestinal epithelial cells. (A)** Images of 3 μm-tissue sections of HT-29-CdtB- and HT-29-H_265_L-derived mice engrafted tumors immunostained for E-cadherin (Adherens junctions) and counterstained with standard hematoxylin staining. HT29-transgenic cell lines were engrafted into immunodeficient mice as previously reported [26]. Magnifications of selected areas are shown in boxes. Black arrows indicate enlarged cells presenting less junctional E-cadherin. Scale bar, 100 µm. **(B)** For MMP gene expression quantification, intestinal epithelial HT-29 and Caco-2 cells were cultured in the presence of doxycycline for 72 h to induce the expression of the CdtB of *H. hepaticus* strain 3B1 or its corresponding inactive form with the H265→L mutation (H_265_L). Then, the relative expression of MMP genes in cells was measured by real-time polymerase chain reaction and normalized according to the expression of the reference gene, hypoxanthine phosphoribosyltransferase 1. Results are the means of 3 independent experiments, each performed in triplicate. Ratios were calculated using the 2^−ΔΔCt^ method. The relative expression rate of MMP genes was reported as a fold increase *versus* H_265_L. The discontinuous line shows the basal rate of MMP expression by H265L cells. **(C)** Wide field images of intestinal epithelial HT-29 and Caco-2 cells cultured in the presence of doxycycline for 72 h to induce the expression of the CdtB of *H. hepaticus* strain 3B1 or its corresponding inactive form with the H_265_→L mutation (H_265_L). Cells were stained with fluorescent-labeled phalloidin (green) to detect F-actin and DAPI to counterstain the nucleus. White arrowheads indicate large lamellipodia. Scale bars, 50 µm for HT-29 and 100 µm for Caco-2. **(D)** Wide field images of intestinal epithelial HT-29 cells cultured in the presence of doxycycline as in S3C. Cells were immunostained with fluorescent antibodies targeting Vimentin or Cortactin (green), and DAPI to counterstain the nucleus (blue). Vimentin and Cortactin expression was quantified and is presented as fold change *versus* H_265_L. The results are presented as the mean ± SD of triplicates in one representative experiment out of three. The discontinuous line shows the basal rate in cells expressing the inactive form of the CdtB of *H. hepaticus* strain, H_265_L. Scale bar, 50 µm. *p<0.05, **p<0.01, ****p<0.0001 *versus* H_265_L. ns, non-significant.

## Notes

### Competing Interest Statement

The authors have declared no competing interest.

